# Vitamin D deficiency and SARS‑CoV‑2 infection: Big-data analysis from March 2020 to March 2021. D-COVID study

**DOI:** 10.1101/2022.10.27.514012

**Authors:** Álvarez M Neira, Jiménez G Navarro, Sánchez N Anguita, Olano M.M Bermejo, R Queipó, Nuñez M Benavent, Jimenez A Parralejo, Yepes G López, Nieto C Sáez

## Abstract

**Background:** Vitamin D has been proposed to have immunomodulatory functions and therefore play a role in coronavirus infection (COVID-19). However, there is no conclusive evidence on its impact on COVID-19 infection and evolution.

**Objective:** To study the association between COVID-19 infection and vitamin D deficiency in patients of a terciary university hospital. To investigate the clinical evolution and prognosis of patients with COVID-19 and vitamin D deficiency.

**Methods:** Using big-data analytics and artificial intelligence through the SAVANA Manager clinical platform, we analysed clinical data from patients with COVID-19 atended in a terciary university hospital from March 2020 to March 2021.

**Results:** Of the 143.157 analysed patients, 36.261 subjects had COVID-19 infection (25.33%); during this period; of these 2588 had vitamin D deficiency (7.14%). Among subjects with COVID-19 and vitamin D deficiency, there was a higher proportion of women OR 1.45 [95% CI 1.33-1.57], adults older than 80 years OR 2.63 [95%CI 2.38-2.91], people living in nursing homes OR 2.88 [95%CI 2.95-3.45] and walking dependence OR 3.45 [95%CI 2.85-4.26]. Regarding clinical course, a higher number of subjects with COVID-19 and vitamin D deficiency required hospitalitation OR 2.41 [95%CI 2.22-2-61], intensive unit care (ICU) OR 2.22 [95% CI 1.64-3.02], had a longer mean hospital stay 3.94 (2.29) p=0.02 and higher mortality OR 1.82 [95%CI 1.66-2.01].)

**Conclusion:** Low serum 25 (OH) Vitamin-D level was significantly associated with a worse clinical evolution and prognosis of COVID-19 infection. We found a higher proportion of institutionalised and dependent people over 80 years of age among patients with COVID-19 and vitamin D deficiency.

## Introduction

Vitamin D is a steroid with functions classically associated with bone metabolism, although in recent years important extraosseous regulatory functions have been described, among which immunomodulatory ones stand out (1). Vitamin D seems to activate innate immunity by stimulating antigen presentation to macrophages, activating neutrophils and T cells and reducing the cytokine storm at the site of infection (2).

In this regard, several hypotheses have been put forward on the role of vitamin D in COVID-19 infection, given that some studies show a higher risk of respiratory infections in patients with vitamin D deficiency and others show a higher incidence of COVID-19 infection and mortality in countries with a high prevalence of vitamin D deficiency (3–7). On the other hand, there is a high prevalence of vitamin D deficiency in the Spanish population and particularly in the elderly and institutionalized population (8–9). Some of the factors associated with vitamin D deficiency are frequently found in this population group: low sun exposure, nutritional disorders or chronic kidney disease. In addition, the confinement imposed in the last two years for disease control may have increased vitamin D deficiency in this group of patients, mainly because of reduced sun exposure and worsening of underlying nutritional disorders.

Therefore, there is a possibility that vitamin D deficiency may be related to a greater severity of symptoms and worse clinical course, with the elderly population being especially vulnerable to this situation.

The main objective of this study is to study the prevalence of vitamin D deficiency in patients with COVID-19 infection admitted to a tertiary hospital during the period March 2020 and March 2021. Secondarily, there are three objectives: 1. to determine the characteristics of patients with vitamin D deficiency and COVID-19 (sociodemographic characteristics, comorbilities, vitamin D treatments, level of walking autonomy, place of residence). 2. Compare clinical course of the disease (need for hospital admission, duration of hospital admission, admission to ICU and mortality) in patients with COVID-19 infection and low or normal vitamin D status, and 3. to study the differences during the three periods of maximum incidence of COVID-19 infection: March 2020 to June 2020 / August 2020 to October 2020 / January 2021 to March 2021.

## Methods

This was a non-interventional, retrospective study using free-text data captured in the EHRs of patients diagnosed with COVID-19 from a terciary Univerity Hospital of Madrid, Spain. The study period was from March 2020 to March 2021. This study was approved by the Ethics and Research Committee of the University Hospital of Getafe (Ref Number: CEIm21/40) on the 31st of September 2021.

Clinical data from a total of 143157 patients were explored. This group of patients fulfils the established inclusion criteria (patients diagnosed with COVID-19 from March 2020 to March 2021) among 2034921 patients with available electronic health records (EHRs) in the Infanta Sofia University Hospital of Madrid Comunity (Spain). Data were collected from all available services, including inpatient and outpatient departments, emergency room and laboratory.

Information from EHRs was extracted using Natural Language Processing (NLP) and artificial intelligence (AI) techniques. SAVANA Manager was used to analysed this free-text clinical information. This software can interpret any content included in EHRs, regardless of the electronic system in which it operates. Importantly, this tool can capture numerical values and clinical notes and transform them into accessible variables, thus allowing for the reuse of the information captured in large-scale collections of clinical records (i.e. big data). The data extraction process has four distinct phases for transferring and aggregating the data into the study database, as follows. 1) Acquisition: data acquisition is the responsibility of the participating site, in close collaboration with SAVANA’s technical staff. The data is pseudonymized using Savana’s “anonymizer” and transferred to Medsavana uploading the information through a Secure File Transfer Protocol (SFTP) available for each center. ; 2) integration: in this phase, data were integrated into the database; 3) NLP: SAVANA’s EHRead technology applied NLP techniques to analyse and extract the unstructured free-text information written in large numbers of EHRs. The NLP output is a structured patient database. EHRead technology creates a completely structured database from unstructured data composed solely of relevant clinical information. This process ensures that personal information is not accessible and makes traceability to individual patients impossible; and 4) validation, consisting of a medical validation of the tool’s output by doctors and researchers.

The terminology used by SAVANA is based on multiple sources such as SNOMED CT [10], which includes medical codes, concepts, synonyms, and definitions regarding symptoms, diagnoses, body structures and substances commonly used in clinical documentation.

Due to the novel methodological approach of this study, we complemented our clinical findings with the assessment of EHRead’s performance. This evaluation was aimed at verifying the system’s accuracy in identifying records that contain mentions of vitamin D deficiency, COVID-19 and its related variables [10]. Briefly, the annotations made by the medical experts were used to generare the gold standard to assess the performance of EHRead’s output; performance is calculated in terms of the standard metrics of accuracy (P), recall (R) and their harmonic mean F-score [11]. The linguistic evaluation of COVID-19 variable has been analyzed in the context of this study, yielding an accuracy, recall and F-score of 0.61, 0.87, and 0.72, respectively. The Interannotator Agreement was 0.70. Other primary variables such as ipovitaminosis D and malnutrition yield and F-score higher tan 0,70.

All statistical analyses were conducted using SPSS software (version 25.0; IBM, Armonk, NY, USA). Unless otherwise indicated, qualitative variables are expressed as absolute frequencies and percentages, while quantitative variables are expressed as mean±SD. For the assessment of statistical significance of numerical variables, we used independent-samples t-tests. To measure the relative distribution of patients assigned to different categories of qualitative variables, we used Chi-squared tests. In all cases, a p-value for statistical significance was set at 0.05.

We followed the Strengthening the Reporting of Observational Studies in Epidemiology (STROBE) guidelines for reporting observational studies [12]. The study was conducted following legal and regulatory requirements and followed research practices described in the International Conference on Harmonisation guidelines for good clinical practice, the Declaration of Helsinki in its latest edition, the guidelines for good pharmacoepidemiology practice and local regulations. Given the retrospective and observational nature of the study, physicians’ prescribing habits and patient assignment to a specific therapeutic strategy were solely determined by the physician, team or hospital concerned. The study involves a combination of structured and unstructured data existing in EHRs and therefore no intervention would be performed on the research subjects. Therefore, the study does not entail any risk for the participants. Likewise, standard informed consent does not apply to this study. The research is with aggregated data and it would be impossible to identify patients in order to seek their informed consent. All actions toward data protection were taken in accordance with the European data protection authorities’ code of good practice regarding big-data projects and the European General Data Protection Regulation (GDPR).

### Study population

Patients from University Infanta Sofia Hospital (Madrid, Spain) with COVID-19 infection diagnosis during the time period from March 2020 to March 2021.

Inclusion criteria: Patients with confirmed diagnosis of COVID-19 infection by PCR technique or antigenic test at Hospital Infanta Sofia in the time period from March 2020 to March 2021.

Patients who do not have the data collected in the clinical history were excluded from the study.

### Study variables

We used the following definitions to categorize study variables:

– COVID-19 infection: considering all those patients with confirmed COVID-19 infection (positive COVID-19 PCR test or antigenic test and associated clinical manifestations).
– Vitamin D deficiency: considering those subjects with a diagnosis of vitamin D deficiency, hypovitaminosis or when vitamin D blood levels were < 30mg/dl.
– Comorbidities: data were collected on the presence of hypertension / diabetes / obesity / malnutrition / dementia / chronic renal failure / liver failure / malabsorption disorders.
– Clinical evolution of Covid 19 infection: the evolution of the patient with COVID 19 infection was collected as: hospital admission / admission to Intensive Care Unit / number of days of hospital admission / mortality.
– Walking independence: information was collected to determine the level of independient ambulation: autonomous / requires technical assistance / total dependen to walk.
– Vitamin D treatments: if the patient was on vitamin D supplementation.
– Time of episode: month of infection

## Results

### Patient characteristics

Of the 143157 analysed patients during the study period (March 2020 to March 2021), 36261 subjects had COVID-19 infection (25.33%) and 2588 of them had also vitamin D deficiency (7.14%).

General characteristics of subjects with COVID-19 and vitamin D deficiency and those without vitamin D deficit are shown in **Table 1**.

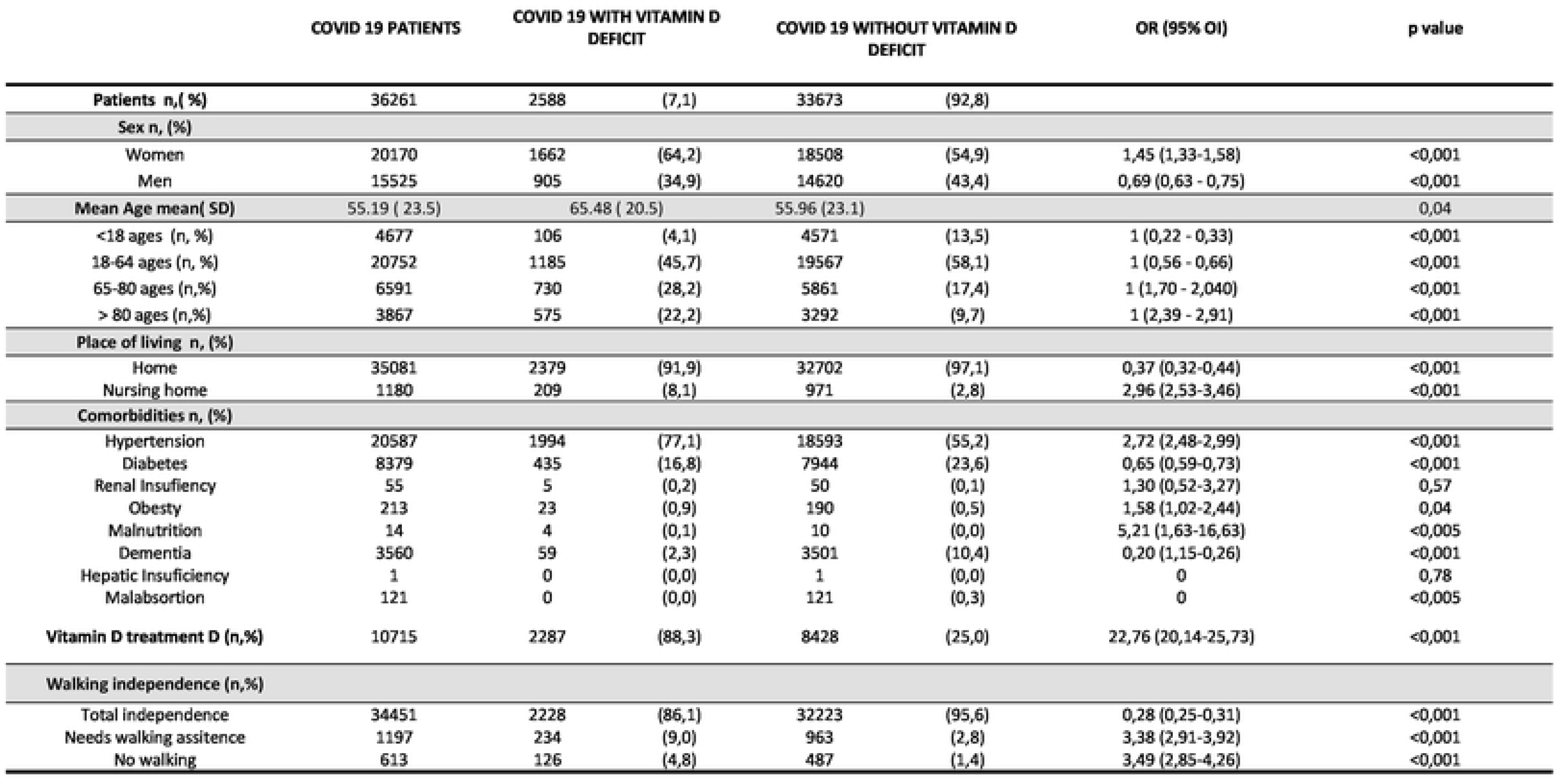

|  | COVID 19 PATIENTS | COVID 19 WITH VITAMIN D DEFICIT |  | COVID 19 WITHOUT VITAMIN D DEFICIT |  | OR (95% OI) | p value |
| --- | --- | --- | --- | --- | --- | --- | --- |
| **Patients n,( %)** | 36261 | 2588 | (7,1) | 33673 | (92,8) |  |  |
| **Sex n, (%)** |  |  |  |  |  |  |  |
| Women | 20170 | 1662 | (64,2) | 18508 | (54,9) | 1,45 (1,33-1,58) | <0,001 |
| Men | 15525 | 905 | (34,9) | 14620 | (43,4) | 0,69 (0,63 - 0,75) | <0,001 |
| **Mean Age mean( SD)** | 55.19 ( 23.5) | 65.48 ( 20.5) |  | 55.96 (23.1) |  |  | 0,04 |
| <18 ages (n, %) | 4677 | 106 | (4,1) | 4571 | (13,5) | 1 (0,22 - 0,33) | <0,001 |
| 18-64 ages (n, %) | 20752 | 1185 | (45,7) | 19567 | (58,1) | 1 (0,56 - 0,66) | <0,001 |
| 65-80 ages (n,%) | 6591 | 730 | (28,2) | 5861 | (17,4) | 1 (1,70 - 2,040) | <0,001 |
| > 80 ages (n,%) | 3867 | 575 | (22,2) | 3292 | (9,7) | 1 (2,39 - 2,91) | <0,001 |
| **Place of living n, (%)** |  |  |  |  |  |  |  |
| Home | 35081 | 2379 | (91,9) | 32702 | (97,1) | 0,37 (0,32-0,44) | <0,001 |
| Nursing home | 1180 | 209 | (8,1) | 971 | (2,8) | 2,96 (2,53-3,46) | <0,001 |
| **Comorbidities n, (%)** |  |  |  |  |  |  |  |
| Hypertension | 20587 | 1994 | (77,1) | 18593 | (55,2) | 2,72 (2,48-2,99) | <0,001 |
| Diabetes | 8379 | 435 | (16,8) | 7944 | (23,6) | 0,65 (0,59-0,73) | <0,001 |
| Renal Insuficiency | 55 | 5 | (0,2) | 50 | (0,1) | 1,30 (0,52-3,27) | 0,57 |
| Obesty | 213 | 23 | (0,9) | 190 | (0,5) | 1,58 (1,02-2,44) | 0,04 |
| Malnutrition | 14 | 4 | (0,1) | 10 | (0,0) | 5,21 (1,63-16,63) | <0,005 |
| Dementia | 3560 | 59 | (2,3) | 3501 | (10,4) | 0,20 (1,15-0,26) | <0,001 |
| Hepatic Insuficiency | 1 | 0 | (0,0) | 1 | (0,0) | 0 | 0,78 |
| Malabsortion | 121 | 0 | (0,0) | 121 | (0,3) | 0 | <0,005 |
| **Vitamin D treatment D (n,%)** | 10715 | 2287 | (88,3) | 8428 | (25,0) | 22,76 (20,14-25,73) | <0,001 |
| **Walking independence (n,%)** |  |  |  |  |  |  |  |
| Total independence | 34451 | 2228 | (86,1) | 32223 | (95,6) | 0,28 (0,25-0,31) | <0,001 |
| Needs walking assitence | 1197 | 234 | (9,0) | 963 | (2,8) | 3,38 (2,91-3,92) | <0,001 |
| No walking | 613 | 126 | (4,8) | 487 | (1,4) | 3,49 (2,85-4,26) | <0,001 |

In COVID-19 with vitamin D deficit there was a higher proportion of women OR 1.45 [95% CI 1.33-1.57] and adults between 65 and 80 years OR 1,86 [95%CI 1,70 - 2,0] and older than 80 years OR 2.63 [95%CI 2.38-2.91].

There was a higher proporción of people living in nursing homes OR 2.88 [95%CI 2.95-3.45] and walking dependence OR 3.45 [95%CI 2.85-4.26].

We found higher proportion of malnutrition, HTA and obesity among patients with COVID-19 and vitamin D deficit and they were more frequently under vitamin D suplements treatments.

### Severity of COVID-19 infection

Regarding clinical course, a higher number of subjects with COVID-19 and vitamin D deficiency required hospitalitation OR 2.41 [95%CI 2.22-2-61] and ICU OR 2.22 [95%CI 1.64-3.02].

Patients with low vitamin D levels had a longer mean hospital stay 3.94 (2.29) p=0.02 and higher mortality OR 1.82 [95%CI 1.66-2.01] (**Table 2**).

**TABLA 2.**
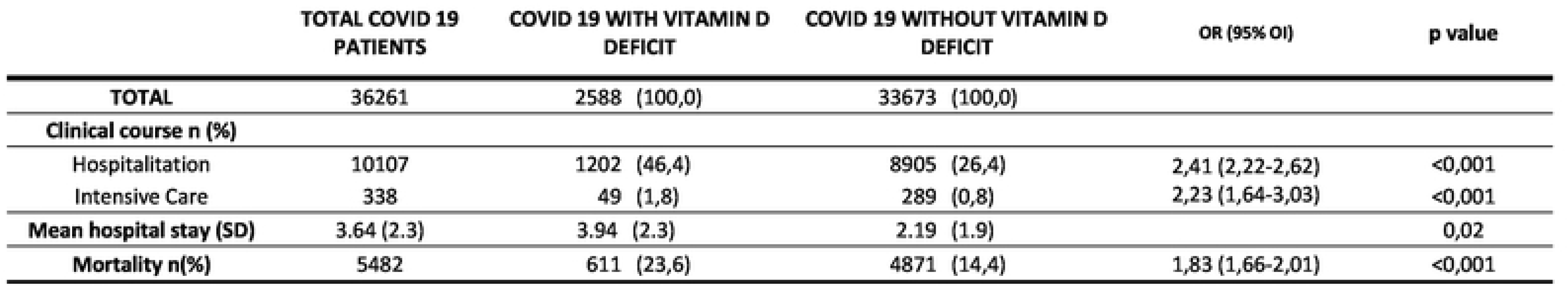
CLINICAL COURSE OF COVID 19 INFECTION.

|  | **TOTAL COVID 19**<br>**PATIENTS** | **COVID 19 WITH VITAMIN D**<br>**DEFICIT** | **COVID 19 WITHOUT VITAMIN D**<br>**DEFICIT** | **OR (95% OI)** | **p value** |
| --- | --- | --- | --- | --- | --- |
| **TOTAL** | 36261 | 2588 (100,0) | 33673 (100,0) |  |  |
| **Clinical course n (%)** |  |  |  |  |  |
| Hospitalitation | 10107 | 1202 (46,4) | 8905 (26,4) | 2,41 (2,22-2,62) | <0,001 |
| Intensive Care | 338 | 49 (1,8) | 289 (0,8) | 2,23 (1,64-3,03) | <0,001 |
| **Mean hospital stay (SD)** | 3.64 (2.3) | 3.94 (2.3) | 2.19 (1.9) |  | 0,02 |
| **Mortality n(%)** | 5482 | 611 (23,6) | 4871 (14,4) | 1,83 (1,66-2,01) | <0,001 |

### Differences during the three periods of maximum incidence

During the first period of pandemia (March to June 2020) patients with COVID-19 and vitamin D deficiency have the same evolution and complications described in the total sample. In the second period (August to October 2020) we find that hospitalitation, Intensive care and mortality are higher in COVID-19 patients with vitamin D deficiency but mean hospital stay is not significatly longer. The third period (January to March 2021) shows differences in clinical course in number of hospitalitations, Intensive care and mortality but not in mean hospital stay (**Table 3**).

**TABLE 3.**
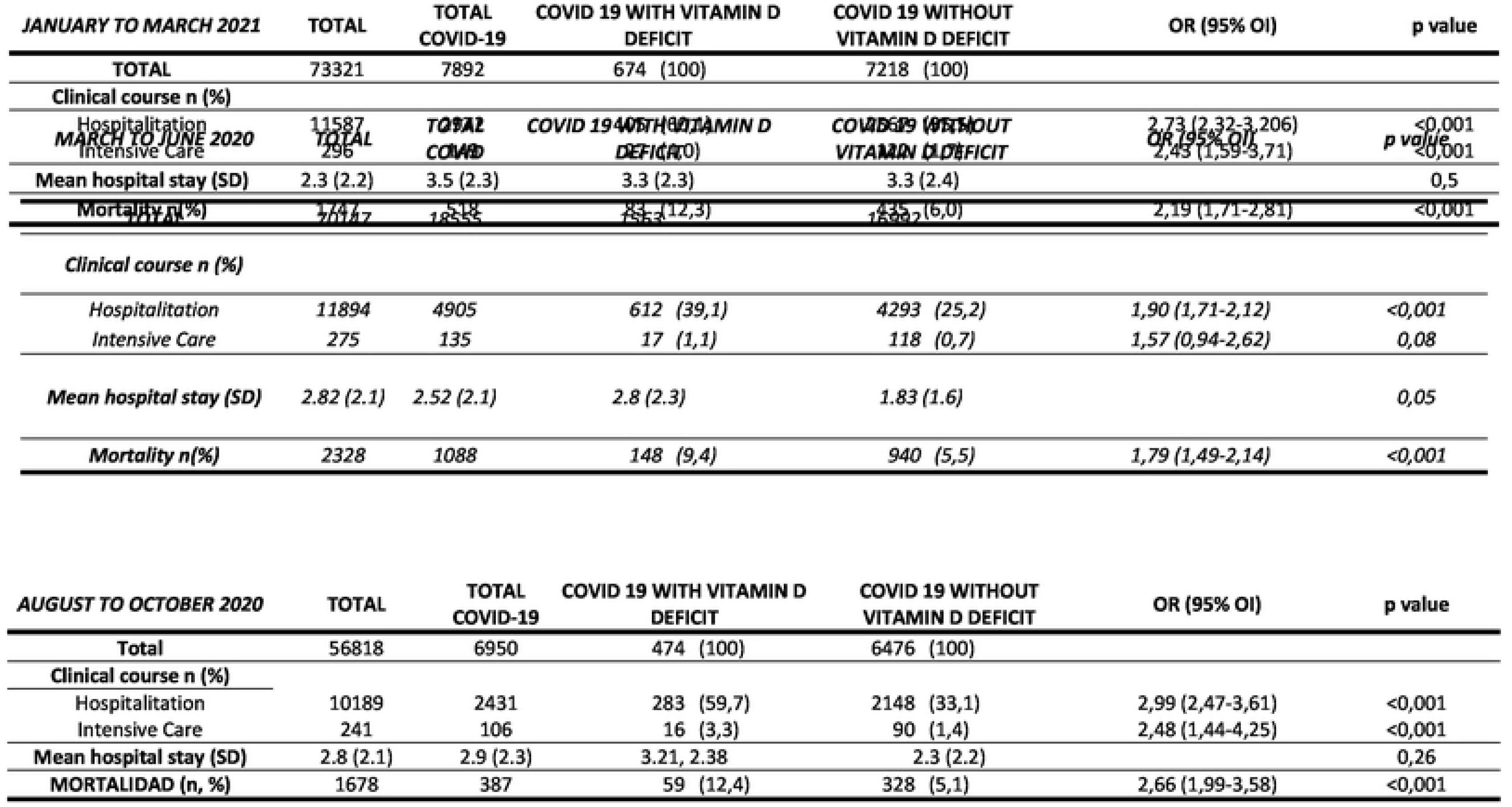
Clinncal course of covid 19 infection during the three periods of máximum incidence.

## Discusión

In this large retrospective case–control study, we study the prevalence, characteristics and evolution of Vitamin D deficit in COVID-19 patients atended in a terciary University Hospital in Madrid (Spain) in the máximum period of pandemia.

We find a significat association between vitamin D deficit and the risk of severe COVID-19 disease when they get infected; higher risk of hospitalitation, more patients require Intensive care attention, higher mortality and longer hospital stay.

Signifcant associations were also found between patients with COVID-19 plus vitamin D deficit and sex, age, place of residence, walking dependence and comorbidity factors. We found no differences between the three periodos of máximum incidence, in number of hospitalitations, Intensive care and mortality but not in mean hospital stay, Our results confrms several others published showing association between low vitamin D levels and COVID-19, particularly those sufering from severe disease and complications [13–15]

Diaz-Curiel et al found similar results in a Spanish population with increased risk of hospital admission and need for critical care, but they didn’t find relationship between vitamin D levels and mortality. [16].

The relationship between vitamin D and COVID-19 infection has been raised because significant association between low serum 25(OH)D levels and the severity of acute respiratory tract infections was found prior to the pandemic [17–18].

Vitamin D modulates angiotensin receptor expression in the lung, stimulates pulmonary surfactant production, reduces hyperinflammatory cytokine storm and increases levels of regulatory T lymphocytes, all of which are closely related to pulmonary infection with RVOC-19 [19].

On the other hand, vitamin D has also important protective efects on the cardiovascular system, including augmentation of myocardial contractility and anti-thrombotic efects and Vitamin D deficit has been associated with other cardiovascular factors such as diabetes, hypertension or dyslipidemia. All these facts could have protective effects against cardiovascular complications in patients with COVID-19 infecction. [20].

Hypovitaminosis D is commonly observed in the elderly population and in this study we find that patients with COVID-19 and vitamin D deficiency are older, with walking dificulties so they depend on others to walk outside, mainly living in nursing homes and with chronic disseases associated to vitamin D deficiency like malnutrition, obesity or real insuficiency.

There are few recent studies studying prevalence of vitamin D deficiency in the Spanish population, although all agree on a high prevalence in this group of age. Gonzalez-Molero et al found that the 33.9% of the Spanish population may be at risk for Vitamin D defict [21] but others describe a prevalence up to 80% in people living in nursing homes. [8–9]. As a result of the high prevalence of vitamin D defict in older adults and the results of this study showing high risk of complications in pacientes with COVID-19 plus vitamin D defict, it sould be positive to consider the benefit of vitamin D suplementation in older population especially those at higher risk (institutionalized or walking dependence).

The main strengths of the present study are the large sample size that has been analyzed and the access to real-world evidence. There are also some limitations; the first one is that, unlike classical research methods, reproducibility is not generally considered in big-data studies, since the latter involves large amounts of information collected from the whole target population. Because we exclusively analysed the data captured in EHRs, the quality of the results reported for some variables is directly tied to the quality of the clinical records; in many cases, EHRs may be partially incomplete and not capture all the relevant clinical information from a given patient. Like in other retrospective studies, some variables can be not properly documented and were therefore not analysed.

Finally, our study sample comprised COVID-19 cases confirmed by PCR test and Antigenic test plus suspected clinical and epidemiological circumstances, so it is posible that some false positive cases could have been included in the final study sample.

## Conclusions

In this large observational population study, we show a signifcant association between vitamin D defciency and the risks of severe disease in patients with COVID-19 infection and a higher proportion of institutionalised and dependent people over 65 years of age among patients with Covid 19 and vitamin D deficiency.

## DECLARATIONS

## Acknowledgements

We thank all the Covid patients because they are the source of the results we explore in this article. We also thank Eduardo Cañada from Computer Dept of Infanta Sofia University Hospital for assitance with analítica codes.

## Ethics approval and consent to participate

The study was conducted according to the guidelines of the Declaration of Helsinki and approved by the Research Ethics Committee of the Hospital Universitario de Getafe (Ref Number: CEIm21/40) on the 31st of September 2021).

## Availability of data and materials

dataset is published in open access in zenodo.

Cited as:

Marta Neira Álvarez, Noemi Anguita Sánchez, Gema Navarro Jiménez, María del Mar Bermejo Olano, Rocío Queipó, María Benavent Nuñez, Alejandro Parralejo Jiménez, Guillermo López Yepes, & Carmen Sáez Nieto. (2022). Vitamin D deficiency and SARS-CoV-2 infection: D-COVID study [Data set].

Zenodo. https://doi.org/10.5281/zenodo.7053208

## Competing interests

Benavent Nuñez M, Parralejo Jimenez A, López Yepes G report that the y are employees at Medsavana. There are no other financial competing interests regarding this paper.

## Funding

There are no non-financial competing interests.

## Authors’ contributions

Conceptualization, M.N.A,

Methodology, investigation, M.N.A, N.J.G, A.S.N, B.O.MM, Q.R, S.N.C, B.N.M, P.J.A, L.Y.G

Writing—Original Draft Preparation, M.N.A, N.J.G, S.N.C

Writing—Review: M.N.A, N.J.G, A.S.N, B.O.MM, Q.R, S.N.C, B.N.M, P.J.A, L.Y.G

All authors have read and agreed to the published version of the manuscript

## Notes

### Competing Interest Statement

The authors have declared no competing interest.

## REFERENCES

1. Charan, J., Goyal, J. P., Saxena, D., & Yadav, P. (2012). Vitamin D for prevention of respiratory tract infections: A systematic review and meta-analysis. In Journal of Pharmacology and Pharmacotherapeutics (Vol. 3, Issue 4, pp. 300–303). doi: 10.4103/0976-500X.103685

2. Sabetta, J. R., Depetrillo, P., Cipriani, R. J., Smardin, J., Burns, L. A., & Landry, M. L. (2010). Serum 25-hydroxyvitamin D and the incidence of acute viral respiratory tract infections in healthy adults. PLoS ONE, 5(6). doi: 10.1371/journal.pone.0011088

3. Laird E, Rhodes J, Kenny RA. Vitamin D and Inflammation: Potential Implications for Severity of Covid-19. Ir Med J. 2020 May 7;113(5):81. PMID: 32603576.

4. Ebadi, M., & Montano-Loza, A. J. (2020). Perspective: improving vitamin D status in the management of COVID-19. In European Journal of Clinical Nutrition (Vol. 74, Issue 6, pp. 856–859). Springer Nature. doi: 10.1038/s41430-020-0661-0

5. Merzon, E., Tworowski, D., Gorohovski, A., Vinker, S., Golan Cohen, A., Green, I., & Frenkel-Morgenstern, M. (2020). Low plasma 25(OH) vitamin D level is associated with increased risk of COVID-19 infection: an Israeli population-based study. FEBS Journal, 287(17), 3693–3702. doi: 10.1111/febs.15495

6. Panagiotou, G., Tee, S. A., Ihsan, Y., Athar, W., Marchitelli, G., Kelly, D., Boot, C. S., Stock, N., Macfarlane, J., Martineau, A. R., Burns, G., & Quinton, R. (2020). Low serum 25-hydroxyvitamin D (25[OH]D) levels in patients hospitalized with COVID-19 are associated with greater disease severity. In Clinical Endocrinology (Vol. 93, Issue 4, pp. 508–511). Blackwell Publishing Ltd. doi:10.1111/cen.14276

7. Lips, P., Cashman, K. D., Lamberg-Allardt, C., Bischoff-Ferrari, H. A., Obermayer-Pietsch, B., Bianchi, M. L., Stepan, J., Fuleihan, G. E. H., & Bouillon, R. (2019). Current Vitamin D status in European and Middle East countries and strategies to prevent Vitamin D deficiency: A position statement of the European Calcified Tissue Society. European Journal of Endocrinology, 180(4), P23–P54. doi: 10.1530/EJE-18-0736

8. Mateo-Pascual, C., Julián-Viñals, R., Alarcón-Alarcón, T., Castell-Alcalá, M. V. ictoria, Iturzaeta-Sánchez, J. M. anuel, & Otero-Piume, A. (2014). Vitamin D deficiency in a cohort over 65 years: prevalence and association with sociodemographic and health factors. Revista Española de Geriatría y Gerontología, 49(5), 210–216. doi: 10.1016/j.regg.2013.11.004

9. Larrosa, M., Gratacòs, J., Vaqueiro, M., Prat, M., Campos Marta Roqué Unidad de Reumatología, F., & Larrosa Padró, D. M. (2021). Valoración del tratamiento sustitutivo. Revista Medicina clínica, 117. doi: 10.1016/S0025-7753(01)72195-1

10. Benson, T. (2010). Principles of Health Interoperability HL7 and SNOMED. ISBN : 978-1-84882-802-5

11. von Elm, E., Altman, D. G., Egger, M., Pocock, S. J., Gøtzsche, P. C., & Vandenbroucke, J. P. (2014). The strengthening the reporting of observational studies in epidemiology (STROBE) statement: Guidelines for reporting observational studies. International Journal of Surgery, 12(12), 1495–1499. doi: 10.1016/j.ijsu.2014.07.013

12. Baeza-Yates, R. A., Ribeiro-Neto, B. Modern Information Retrieval. Boston, Addison-Wesley Longman Publishing Co, 1999. ISBN:978-0-201-39829-8

13. Israel, A., Cicurel, A., Feldhamer, I., Stern, F., Dror, Y., Giveon, S. M., Gillis, D., Strich, D., & Lavie, G. (2022). Vitamin D deficiency is associated with higher risks for SARS-CoV-2 infection and COVID-19 severity: a retrospective case–control study. Internal and Emergency Medicine, 17(4), 1053–1063. doi: 10.1007/s11739-021-02902-w

14. Crafa A, Cannarella R, Condorelli RA, Mongioì LM, Barbagallo F, Aversa A, La Vignera S, Calogero AE. Influence of 25-hydroxy-cholecalciferol levels on SARS-CoV-2 infection and COVID-19 severity: A systematic review and meta-analysis. EClinicalMedicine. 2021 Jul;37:100967. doi: 10.1016/j.eclinm.2021.100967. Epub 2021 Jun 18. Erratum in: EClinicalMedicine. 2021 Nov;41:101168. PMID: 34179737; PMCID: PMC8215557.

15. Ye, K., Tang, F., Liao, X., Shaw, B. A., Deng, M., Huang, G., Qin, Z., Peng, X., Xiao, H., Chen, C., Liu, X., Ning, L., Wang, B., Tang, N., Li, M., Xu, F., Lin, S., & Yang, J. (2021). Does Serum Vitamin D Level Affect COVID-19 Infection and Its Severity?-A Case-Control Study. Journal of the American College of Nutrition, 40(8), 724–731. doi: 10.1080/07315724.2020.1826005

16. Diaz-Curiel, M., Cabello, A., Arboiro-Pinel, R., Mansur, L., Heili-Frades, S., Mahillo-Fernandez, I., Herrero-González, A., & Andrade-Poveda, M. (2021). The relationship between 25(OH) vitamin D levels and COVID-19 onset and disease course in Spanish patients. Journal of Steroid Biochemistry and Molecular Biology, 212. doi: 10.1016/j.jsbmb.2021.105928

17. Holick, M. F. (2007). Medical Progress Vitamin D Deficiency. In N Engl J Med (Vol. 357). www.nejm.org

18. Sabetta, J. R., Depetrillo, P., Cipriani, R. J., Smardin, J., Burns, L. A., & Landry, M. L. (2010). Serum 25-hydroxyvitamin D and the incidence of acute viral respiratory tract infections in healthy adults. PLoS ONE, 5(6). doi: 10.1371/journal.pone.0011088

19. Xu, Y., Baylink, D. J., Chen, C. S., Reeves, M. E., Xiao, J., Lacy, C., Lau, E., & Cao, H. (2020). The importance of Vitamin D metabolism as a potential prophylactic, immunoregulatory and neuroprotective treatment for COVID-19. In Journal of Translational Medicine (Vol. 18, Issue 1). BioMed Central. doi: 10.1186/s12967-020-02488-5

20. Driggin, E., Madhavan, M. v., & Gupta, A. (2022). The role of vitamin D in cardiovascular disease and COVID-19. In Reviews in Endocrine and Metabolic Disorders (Vol. 23, Issue 2, pp. 293–297). Springer. doi: 10.1007/s11154-021-09674-w

21. González-Molero, I., Morcillo, S., Valdes, S., Pérez Valero, V., Botas, P., Delgado, E., Hernandez, D., Olveira, G., Rojo-Martinez, G., Gutierrez, C., Valero, P., Repiso, G. C., & Martin, R. E. (2010). Vitamin D deficiency in Spain: a population-based cohort study 1 Vitamin D deficiency in Spain: a population-based cohort study. European Journal of Clinical Nutrition. doi: 10.1038/ejcn.2010.265ï

